# Physical and functional interaction of Lrrc56 and Odad3 controls deployment of axonemal dyneins in vertebrate multiciliated cells

**DOI:** 10.1101/2025.07.02.662827

**Authors:** Nayeli G. Reyes-Nava, Chanjae Lee, Ophelia Papoulas, Juyeon Hong, Edward M. Marcotte, John B. Wallingford

## Abstract

Primary ciliary dyskinesia is a genetically heterogeneous motile ciliopathy characterized chronic respiratory disease, laterality defects, hydrocephalus, and infertility, caused by impaired function of motile cilia. *LRRC56* has recently emerged as a novel PCD candidate gene, but its role in vertebrate cilia remains poorly understood. Here, we used *Xenopus laevis* multiciliated cells, targeted knockdown, and *in vivo* imaging to investigate *Lrrc56* function. We show that loss of *lrrc56* causes specific depletion of outer dynein arms (ODAs) from the distal axoneme. *In vivo* affinity purification mass spectrometry revealed that Lrrc56 binds the ODA docking complex components, including Odad3. Consistently, *Lrrc56* knockdown also led to distal loss of Odad3. Moreover, we show that disease-associated variants in *LRRC56* and *ODAD3* disrupted their localization and interaction, pointing to a shared functional pathway. Our work demonstrates that *Lrrc56* is a critical regulator of distal ODAs and ODA docking complex deployment and provides new mechanistic insight into how *LRRC56* mutations contribute to PCD.

## Introduction

Motile ciliopathies are a group of inherited disorders caused by structural and/or functional defects in motile cilia, microtubule-based organelles that generate directional fluid flow across epithelial surfaces. These disorders affect multiple organ systems and are commonly associated with chronic respiratory infections, laterality defects, hydrocephalus, and infertility (Wallmeier et al., 2020). Among them, primary ciliary dyskinesia (PCD) is the most extensively characterized at the genetic level, with more than 40 causative genes identified to date (Horani & Ferkol, 2021; Raidt et al., 2023). However, the clinical manifestations of PCD are heterogeneous and often overlap with other respiratory conditions, complicating diagnosis. Despite growing insights into the genetic underpinnings, many PCD-associated genes remain poorly characterized, and key molecular mechanisms underlying motile cilia dysfunction are still not fully understood.

Motile cilia beating is driven by axonemal dynein complexes like outer dynein arms (ODAs), which line along the microtubule doublets to generate ciliary motion (King, 2016). ODA motors are preassembled in the cytoplasm (Fowkes & Mitchell, 1998) through the coordinated action of dynein axonemal assembly factors (DNAAFs) and chaperones (King, 2018; Qiu & Roy, 2022). ODA preassembly takes within specialized, membraneless organelles known as DynAPs (dynein axonemal particles), where DNAAFs, along with other assembly machinery, localize (Huizar et al., 2018) (Fig. 1A).

**Figure 1.**
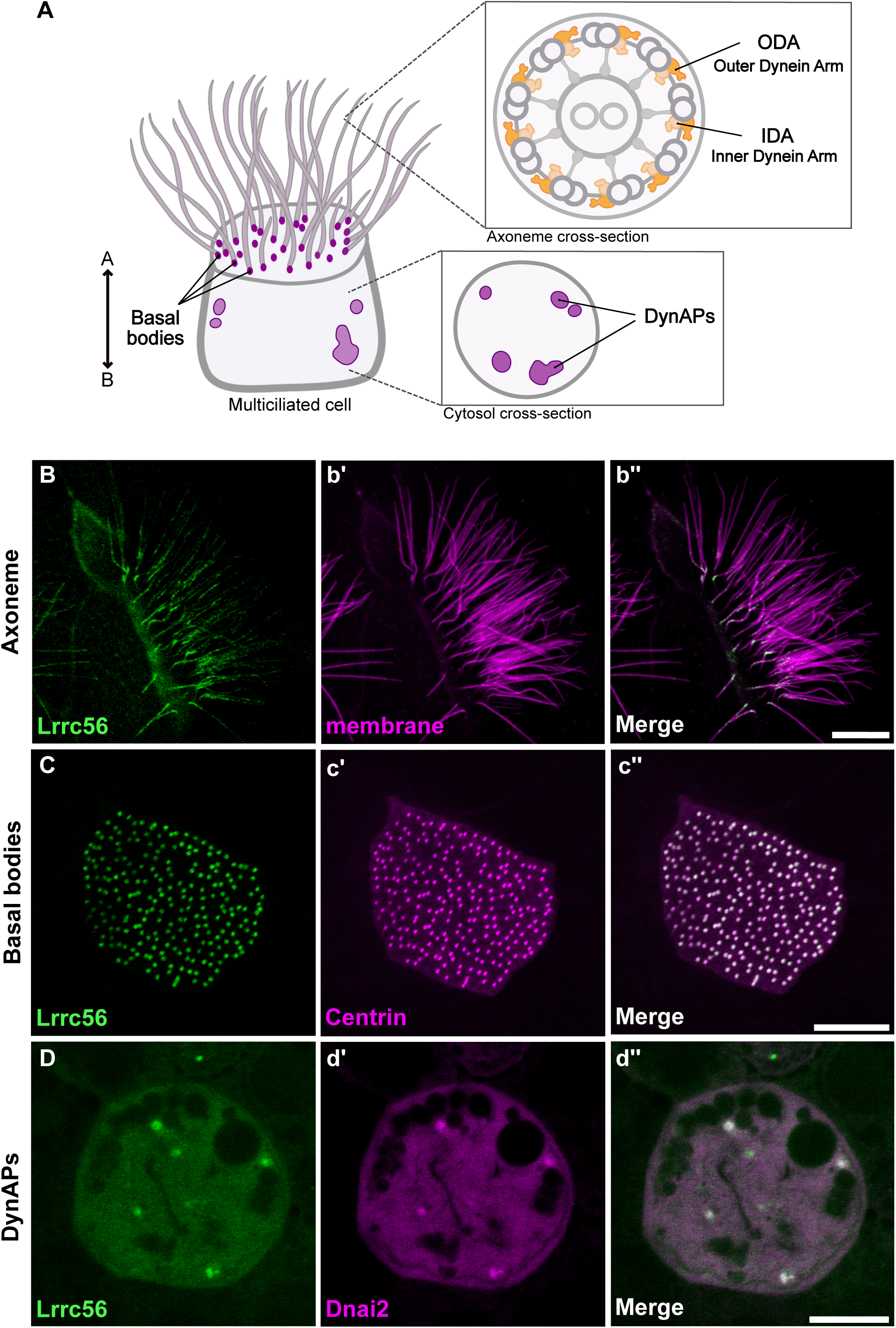
Localization of GFP-Lrrc56 in *Xenopus* MCCs. (A) Schematic of a vertebrate multiciliated cell (MCC). The upper insert shows a cross-section of an axoneme with the relative positions of outer dynein arms (ODA) and inner dynein arms (IDA). The bottom insert illustrates a representative cross-section of the MCC cytoplasm, highlighting DynAPs. (B-D) Representative *in vivo* confocal images showing GFP-Lrrc56 (green) localization along the length of ciliary axonemes, basal bodies, and at cytosolic foci (DynAPs) in *Xenopus* MCCs. (b’-d’) Membrane labeling with CAAX-RFP (magenta) marks motile cilia, centrin-RFP (magenta) marks basal bodies, and mcherry-Dnai2 (magenta) labels DynAPs. (b’’-d’’) Representative merged confocal images from the previous channels showing GFP-Lrrc56 (green) and the various markers (magenta). Scale bars = 10 µm.

Following assembly, ODAs must be selectively and efficiently transported from DynAPs through the cytoplasm to the base of the cilium. This trafficking process is thought to involve specific transport adaptors and cytoplasmic motor proteins that mediate the handoff of ODAs to the intraflagellar transport (IFT) machinery (Hibbard et al., 2022; Lechtreck, 2022). Once inside the cilium, ODAs are anchored to axonemal microtubules via the ODA docking complex (Gui et al., 2021). Critically, mutations in dynein subunits, DNAAFs, or ODA docking complex components disrupt this assembly and trafficking pipeline, resulting in motile ciliopathies (Bonnefoy et al., 2018; Leslie et al., 2022; Qiu & Roy, 2022). While a few molecular players have been identified, the precise mechanisms by which ODAs are transported from DynAPs to the ciliary base remain poorly understood. Because proteins involved in dynein trafficking are frequently implicated in human disease, they present an entry point for understanding the mechanistic basis of these diseases.

One such protein, LRRC56, is a leucine-rich repeat protein whose role in ODA transport and assembly was first proposed based on studies of its algal homolog ODA8 in *Chlamydomonas*. Mutations in *LRRC56* have been associated with chronic respiratory disease and laterality defects in humans, consistent with the clinical presentation of PCD (Alasmari et al., 2022; Asseri et al., 2023; Bonnefoy et al., 2018). Surprisingly, despite these clinical phenotypes, transmission electron microscopy (TEM) of patient samples shows apparently intact ODAs. In *Trypanosoma*, loss of Lrrc56 disrupts ciliary motility and leads to selective loss of ODAs at the distal axoneme (Bonnefoy et al., 2018). More recently, murine studies show that Lrrc56 deficiency results in ciliary ultrastructural abnormalities and phenotypes consistent with PCD (Wu et al., 2025). Together, these findings highlight the clinical significance of *LRRC56* and the variability in ciliary ultrastructural phenotypes across species. Although work in unicellular models points to a role in ODA trafficking and distal targeting, how Lrrc56 functions in vertebrate cilia assembly and contributes to human disease remains an open and important question.

Here, we characterize Lrrc56 in vertebrate multiciliated cells (MCCs) and demonstrate its essential role in ODA deployment. In *Xenopus*, Lrrc56 localizes to DynAPs, basal bodies, and axonemes. Importantly, ciliopathy-associated alleles in *LRRC56* disrupt its normal subcellular localization. Specifically, mutations in the leucine-rich repeat (LRR) domains and deletions of the intrinsically disordered regions (IDRs) result in distinct phenotypes, suggesting domain-specific functions for Lrrc56. Moreover, variants in *ODAD3*—an ODA docking complex component identified here as a binding partner of Lrrc56—not only alter its localization but also disrupt its functional interaction with Lrrc56. Together, these findings define a conserved role for Lrrc56 in vertebrate MCCs and provide mechanistic insight into the pathogenesis of *LRRC56*-related motile ciliopathies.

## Results

### Lrrc56 localization pattern suggests a transport function

In ciliated unicellular organisms, like *Chlamydomonas* and *Trypanosoma*, Lrrc56 orthologues localize to both the flagella and cell body (Bonnefoy et al., 2018; Desai et al., 2015); however, its localization in vertebrate MCCs has yet to be described. We therefore turned to *Xenopus* embryos MCCs, which are highly amenable to live imaging and a reliable model of the biology of mammalian MCCs (Walentek & Quigley, 2017).

We found that GFP-Lrrc56 localized to motile cilia in *Xenopus* MCCs, displaying a punctate distribution along the axoneme with slight enrichment at the proximal end (Fig. 1B,b’). Additionally, Lrrc56 was strongly enriched at basal bodies, co-localizing with Centrin-BFP (Fig. 1C,c’), a pattern not previously reported in unicellular organisms.

ODAs are thought to be preassembled in DynAPs before being transported to cilia (Huizar et al., 2018), a process in which Lrrc56 has been implicated (Desai et al., 2015). We were interested, then to observe that Lrrc56 was also enriched in foci in the cytoplasm (Fig. 1D). We confirmed that these foci reflect the ODA-specific region of DynAps (Lee et al., 2020) by co-labeling with the ODA subunit mCherry-Dnai2 (Fig. 1D’). Thus, Lrrc56 localizes to DynAPs, basal bodies, and axonemes, consistent with its proposed role in trafficking of ODAs.

### *LRRC56* pathogenic variants associated with PCD disrupt its localization to MCCs

Pathogenic mutations in *LRRC56* have been implicated in PCD (Fig. 2A) (Alasmari et al., 2022; Asseri et al., 2023; Bonnefoy et al., 2018). While one of these variants was linked to cilia motility in trypanosomes (Bonnefoy et al., 2018), the effect of these alleles in vertebrate MCCs is unknown. These disease variants affect residues conserved across species (Fig. S1), allowing precise modeling of the corresponding mutations in the *Xenopus* ortholog (Fig. 2A). To test whether ciliopathy alleles alter compartment-specific distribution, we generated the equivalent alleles in *Xenopus* Lrrc56 and expressed them in MCCs.

**Figure 2.**
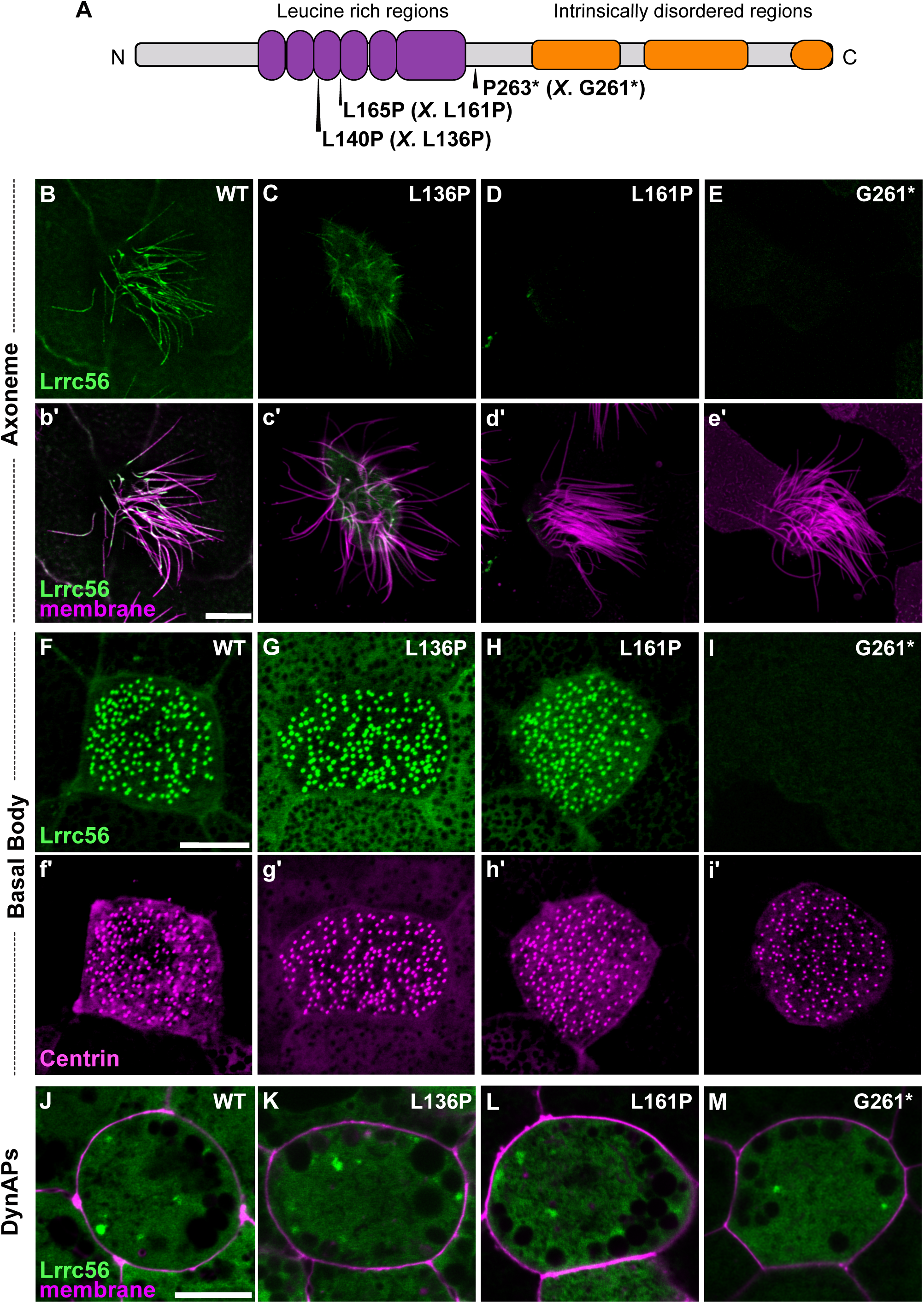
Lrrc56 PCD variants show a spectrum of localization defects. (A) Schematic of LRRC56 (UniProt ID: Q8IYG6), showing the leucine-rich regions and long intrinsically disordered C-terminus. The location of three human motile ciliopathy-causing variants L140P, L165P, and D266* is indicated. Corresponding *Xenopus* allele is in parenthesis and indicated by an *X.* (B-E) En face images of single multiciliated cells (MCCs) showing Lrrc56 (green) axoneme localization for wild-type (WT), L136P (human L140P), L161P (human L165P), and D261* (human P263*) variants. Lrrc56 axonemal localization is disrupted in all three variants. (b’-e’) Merged images showing Lrrc56 (green) with membrane labeling (magenta) reveals normal motile cilia structure. (F-I) Representative *en face* images of Lrrc56 basal body localization for WT, L136P, L161P, and G261* variants. Variants located at leucine rich domains show normal localization to basal bodies, but G261* variant fails to localize at basal bodies. (f’-i’) Centrin (magenta) labeling of basal bodies for the corresponding WT, L136P, L161P, and G261* variants. (J-M) Merged channels showing Lrrc56 (green) with membrane (magenta) reveal normal localization at DynAPs for WT, L136P, L161P, and G261* variants. Scale bars = 10 µm.

The L140P variant changes a well conserved residue corresponding to *Xenopus* L136P within the leucine-rich repeat (LRR) domain of Lrrc56 (Fig. 2A; Fig. S1). This variant showed reduced axonemal enrichment (Fig. 2C), while still localizing to basal bodies and DynAPs (Fig. 2G, K). Notably, this subcellular mislocalization contrasts with reports in *Trypanosoma*, where the equivalent mutation did not alter Lrrc56 localization (Bonnefoy et al., 2018), highlighting potential species-specific differences in localization or function.

L165P—the most recurrent *LRRC56* variant found in PCD patients (Alasmari et al., 2022; Asseri et al., 2023)– is also well conserved (Fig. S1) and the corresponding *Xenopus* L161P allele showed a complete loss of axonemal localization (Fig. 2D). Like L136P, this allele still accumulated at basal bodies and in DynAPs (Fig. 2H, L). Taken together, these two alleles suggest a requirement for the LRR domain in directing Lrrc56 to motile cilia.

Finally, we examined the P263* nonsense mutation (corresponding to *Xenopus* G261*), which truncates the C-terminal intrinsically disordered region (IDR) of Lrrc56 (Fig. 2A; Fig. S1) Notably, loss of the IDR completely abolished Lrrc56 localization to both axonemes and basal bodies (Fig. 2E, I). However, the G261* variant remained enriched at DynAPs (Fig. 2M). Western blot analysis confirmed that all variants were well-expressed (Fig. S1), indicating that mislocalization is not due to protein instability, but rather reflects a defect in trafficking.

Together, these findings reveal that pathogenic *LRRC56* variants disrupt Lrrc56 trafficking to motile cilia in distinct ways, suggesting that different domains of the protein contribute uniquely to its localization and function. While the LRR domains appear essential for axonemal targeting, the IDRs may play roles in cytosolic trafficking or basal body association. This work establishes a framework for dissecting the compartment-specific regulation of Lrrc56—and how its mislocalization may contribute to PCD pathogenesis.

### Lrrc56 is essential for ODA deployment to motile cilia in *Xenopus*

Partial or complete loss of ODAs have been reported in unicellular organisms lacking Lrrc56 whereas *LRRC56*-mutant human respiratory cells show apparently normal axonemal ODA ultrastructure via TEM (Bonnefoy et al., 2018; Kamiya, 1988). Thus, to further explore the role of Lrrc56 in ODA deployment to axonemes in vertebrates, we performed loss of function analysis using knockdown (KD) of Lrrc56. An antisense morpholino oligonucleotide designed to block Lrrc56 splicing severely depleted the mRNA as indicated by reverse transcription PCR (RT-PCR) (Fig. S3).

Lrrc56 knockdown disrupted the normal axonemal distribution of the ODA light chain subunit Dnal4 (GFP-Dnal4), which is typically enriched along the length of the axoneme (Fig. 3A, B). This defect was rescued by re-expression of Flag-tagged Lrrc56 (Fig. 3C), demonstrating the specificity of the KD. Quantification—normalized to a membrane-RFP marker—confirmed a significant reduction in Dnal4 levels in knockdown MCCs that was restored upon rescue (Fig. 3D). To confirm this result, we examined an additional ODA subunit, the intermediate ODA subunit Dnai2. We observed a similar phenotype, with Lrrc56 knockdown decreasing GFP-Dnai2 axonemal signal and rescue restoring it (Fig. 3E–H).

**Figure 3.**
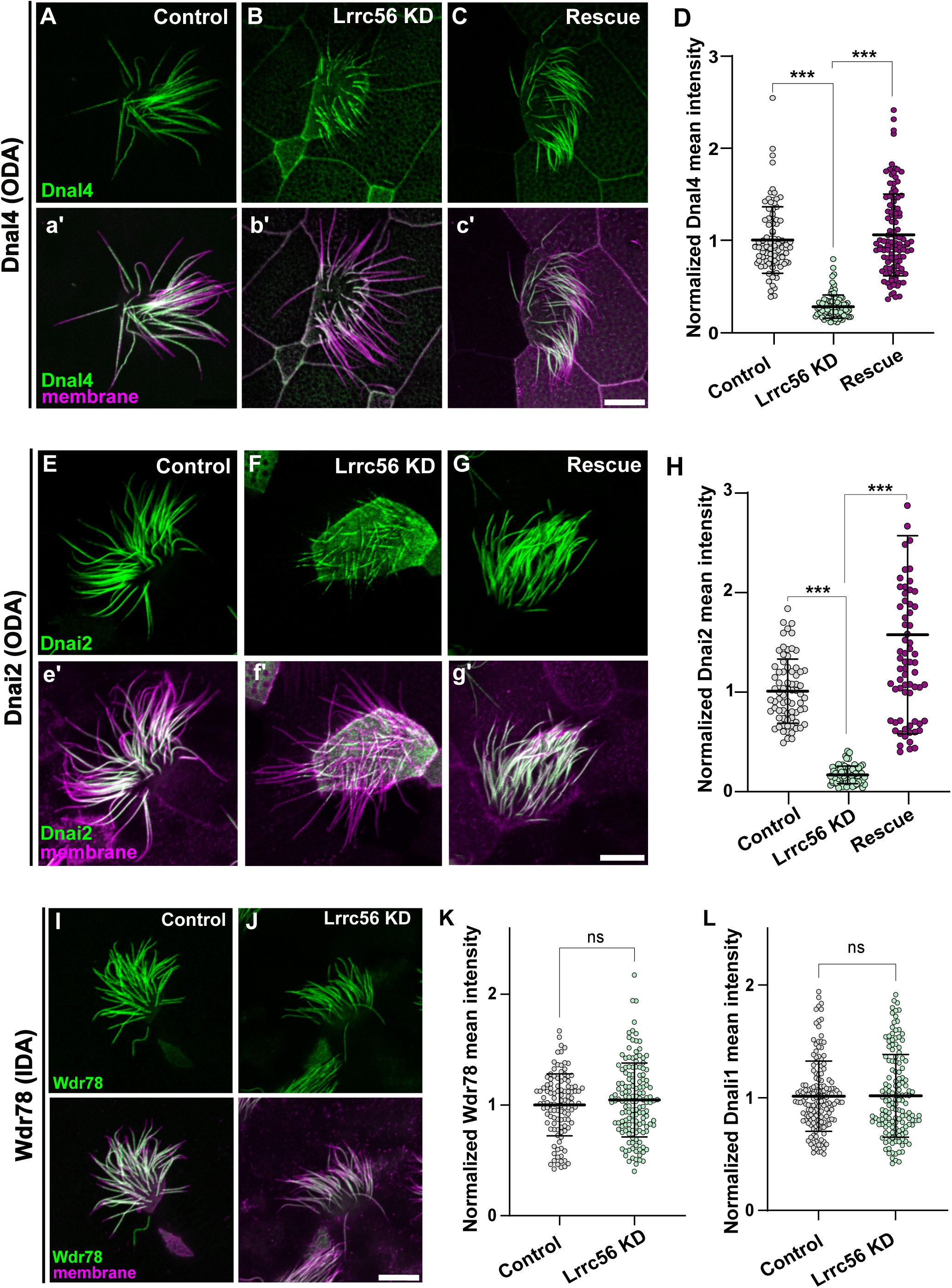
Lrrc56 is essential for ODA deployment to axonemes. (A) GFP-Dnal4 labeling of motile axonemes in control MCCs shows normal localization of outer dynein arm (ODA) subunits. (a’) Merged channels (membrane labeled with CAAX-RFP, magenta, and Dnal4, green) reveal normal cilia morphology in control MCCs. (B) GFP-Dnal4 is absent from motile cilia in Lrrc56 knockdown (Lrrc56 KD) MCCs. (b’) Membrane labeling (magenta) in merged channels shows normal motile cilia morphology despite Lrrc56 KD. (C) GFP-Dnal4 is restored to Lrrc56 KD motile cilia after ectopic expression of FLAG-Lrrc56. (c’) Merged channels show membrane (magenta) and Dnal4 (green) labeling in rescued MCCs. (D) Quantification of mean ± standard deviation of normalized GFP-Dnal4 fluorescence (see methods) in axonemes of control (n=91), Lrrc56 KD (n=97), and rescue (n=11) MCCs. N > 25 cells from 9 embryos across 3 experiments for all conditions. (E-G) GFP-Dnai2 labeling of motile axonemes in Control, Lrrc56 KD, and rescue MCCs. GFP-Dnai2 shows normal localization in control MCCs but is lost from axonemes in Lrrc56 KD MCCs. Loss of GFP-Dnai2 is restored after ectopic expression of Lrrc56. (e-g) Merged channels show membrane (magenta) and Dnai2 (green) labeling for cells in panels E-G. (H) Quantification of normalized GFP-Dnai2 mean intensity along axonemes in control (n=73), Lrrc56 KD (n=73), and rescue (n=74) MCCs. N > 25 cells from 9 embryos across 3 experiments for all conditions. (I) GFP-Wdr78 labeling of inner dynein arm (IDA) subunits shows normal localization in motile axonemes. (J) GFP-Wdr78 remains localized to motile cilia in Lrrc56 KD MCCs. (J’) Merged channels show normal cilia morphology in both control and Lrrc56 KD MCCs, with membrane (magenta) and Wdr78 (green) labeling. (K-L) Graphs showing mean intensity of normalized IDA subunits Wdr78 (K) and Dnali4 (L) in control and Lrrc56 KD MCCs. Scale bars = 10 µm.

In human patients, inner dynein arms (IDAs) were unaffected after loss of LRRC56, and the same was true in unicellular organisms (Bonnefoy et al., 2018; Desai et al., 2015). Accordingly, we found that Lrrc56 knockdown did not disrupt the axonemal localization of the IDA subunits GFP-Wdr78 and GFP-Dnali1 (Fig. 3I–L). These findings support the conclusion that Lrrc56 is specifically required for deployment to axonemes of specific subunits of the ODA, but not IDA, in *Xenopus* MCCs.

### *In vivo* APMS reveals interaction between Lrrc56 and docking complex proteins

Although LRRC56 is linked to PCD, its molecular function remains unknown. We therefore sought to define the interaction landscape of Lrrc56 specifically in MCCs using *in vivo* affinity purification followed by mass spectrometry (AP-MS). We expressed GFP-tagged Lrrc56 under the control of an MCC-specific *α*-tubulin promoter (Deblandre et al., 1999) and dissected ∼750 ectodermal explants (“animal caps”) from *Xenopus* embryos. Upon culture, these explants differentiated into mucociliary epithelium containing beating MCCs (Drysdale & Elinson, 1992; Walentek & Quigley, 2017). Protein was isolated and GFP-tagged Lrrc56 was immunoprecipitated using anti-GFP agarose beads. We performed the same method with GFP alone and subtracted the resulting background to calculate fold enrichment for Lrrc56-specific interactors (see Methods; Fig. S4).

As expected, the bait protein Lrrc56 itself was the most strongly enriched hit, serving as a robust positive control (Fig. 4A). Moreover, our AP-MS analysis revealed strong enrichment of two key ODA docking complex components: Odad1 (Ccdc114) and Odad3 (Ccdc151) (Fig. 4A). These coiled-coil proteins are central to the pentameric docking complex, recently resolved by cryo-EM of bovine respiratory cilia (Gui et al., 2021). The complex—composed of Odad1, Odad2 (Armc4), Odad3, Odad4 (Ttc25), and Odad5 (Calaxin)—forms a crucial bridge anchoring ODAs to the axonemal microtubule doublets (Gui et al., 2021; Yamaguchi et al., 2023). Importantly, mutations in *ODAD3* and *ODAD1* in humans are associated with motile ciliopathies, including PCD (Alsaadi et al., 2014; Hjeij et al., 2014; Knowles et al., 2013; Li et al., 2019). Co-enrichment of Odad1 and Odad3 with Lrrc56 strongly suggests a functional link between Lrrc56 and ODA docking machinery.

We therefore explored these interactions using protein structure modeling (see Methods; Fig. 4B). Initial AlphaFold3 modeling of full-length Lrrc56, Odad3, and Odad1 yielded a low-confidence multimer (ipTM = 0.41; pTM = 0.38), but the model had regions of high per-residue confidence scores (pIDDT > 70) for the Lrrc56 LRR domain and the associated coiled-coil regions of Odad3 and Odad1. These regions (residues 42–261 of Lrrc56, 162–239 of Odad3, and 137– 220 of Odad1) were subsequently modeled in isolation, resulting in a higher-confidence structure (ipTM = 0.74; pTM = 0.69) (Fig. 4B). Consistent with these values, the predicted aligned error (PAE)—a measure of AlphaFold3’s confidence in the relative positioning of residue pairs— indicated a model with low global alignment error (Fig. 4D).

The ciliopathy-associated alleles L136P and L161P mapped to the core LRR domain of Lrrc56, positioned immediately adjacent to the predicted Odad3 interaction surface (Fig. 4E). These substitutions are likely to disrupt LRR folding, contributing to the loss of axonemal localization observed experimentally. In contrast, the G261* truncation lies at the C-terminal end of the modeled region and removes the unstructured IDRs (not included in the structure), suggesting that these distal regions are critical for proper localization and potentially for interactions beyond the modeled core (Fig. 4E). Human LRRC56, ODAD3, and ODAD1 models were also generated and show similar structures/folding as well as localization of ciliopathy associated alleles (Fig. S5).

**Figure 4.**
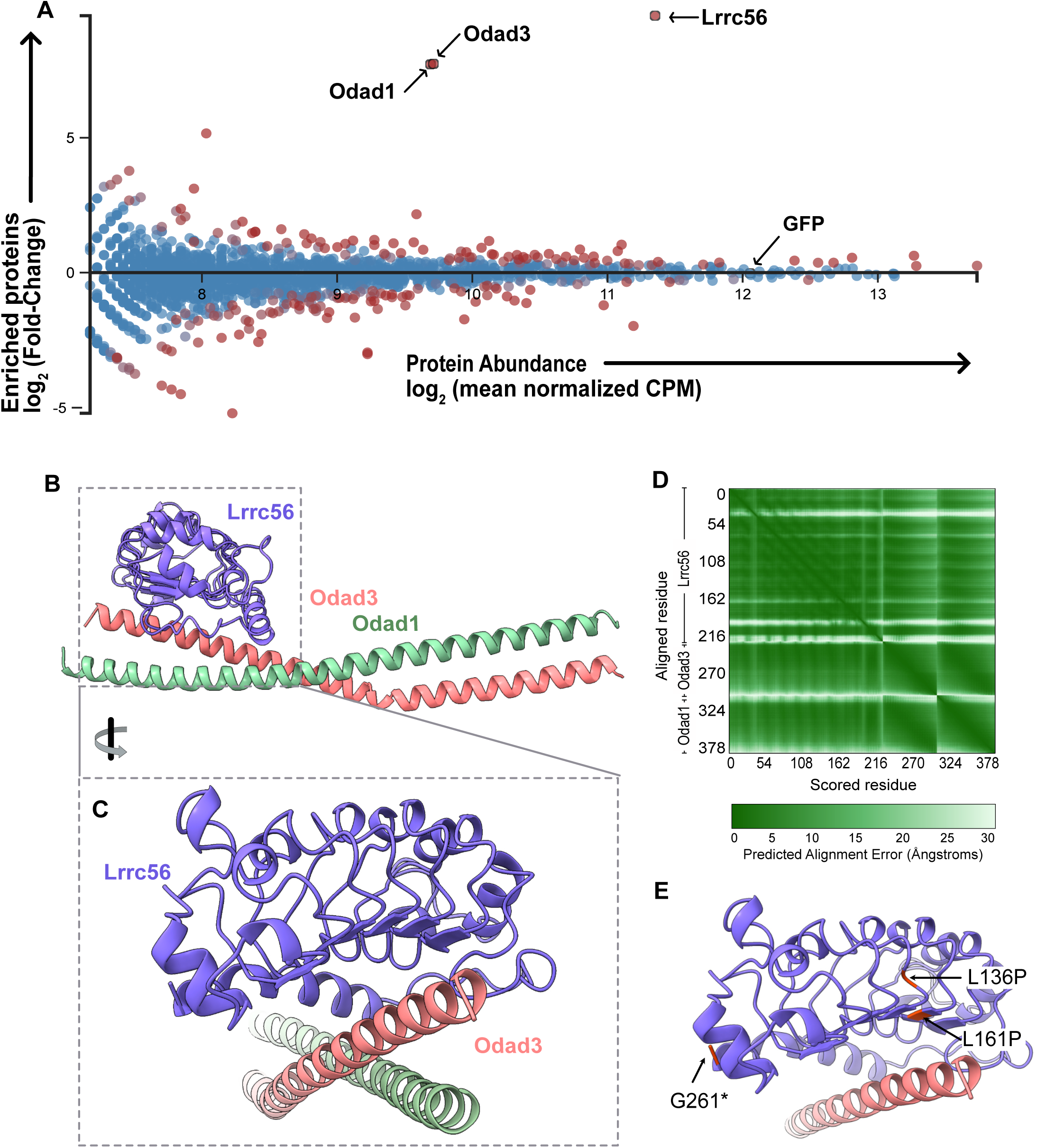
*In Vivo* Affinity-Purification Mass-spectrometry (AP-MS) reveals novel interactors of Lrrc56. (A) MA plot of enriched proteins (log2 fold change) and average abundance between GFP-Lrrc56 (experiment) and GFP (control) immunoprecipitated proteins. Top hits are indicated with lack arrows. Lrrc56 (bait) is highly enriched and abundant in the experiment group. Odad1 and Odad3 are the top two enriched proteins in Lrrc56 IP group. GFP remains unchanged between control and experiment groups. (B) AlphaFold3-predicted structure of *Xenopus* Lrrc56 and its interactors, Odad3 and Odad1. Each monomer is color-coded as indicated. The model includes residues 42–261 of Lrrc56, 162–239 of Odad3, and 137–220 of Odad1. (C) Enlarged view of the Lrrc56–Odad3 interface. (D) Predicted Aligned Error (PAE) plot for the interaction between *Xenopus* Lrrc56, Odad3, and Odad1 in the model shown in (B). (E) Close-up of the Lrrc56–Odad3 interface in the AlphaFold3 model, highlighting the positions of Lrrc56 ciliopathy-associated variants (L136P, L161P, and G261*).

### Lrrc56 is necessary for deployment of Odad3 to the distal end of the axoneme

Because AlphaFold predicted a close interaction between Lrrc56 and Odad3, we asked whether disruption of Lrrc56 would affect Odad3 localization. In control MCCs Odad3 localized along the length of the axoneme, excluding the most distal region, which is known to be enriched in Spef1 (Fig. 5A, a’). However, following Lrrc56 knockdown, the Odad3-GFP signal was significantly reduced along the axoneme compared to controls (Fig. 5B). To quantify this change, the mean intensity of Odad3-GFP along the axoneme was measured and normalized to a membrane marker. As shown in Figure 5C, the mean Odad3-GFP intensity in Lrrc56-depleted MCCs was significantly decreased relative to controls.

**Figure 5.**
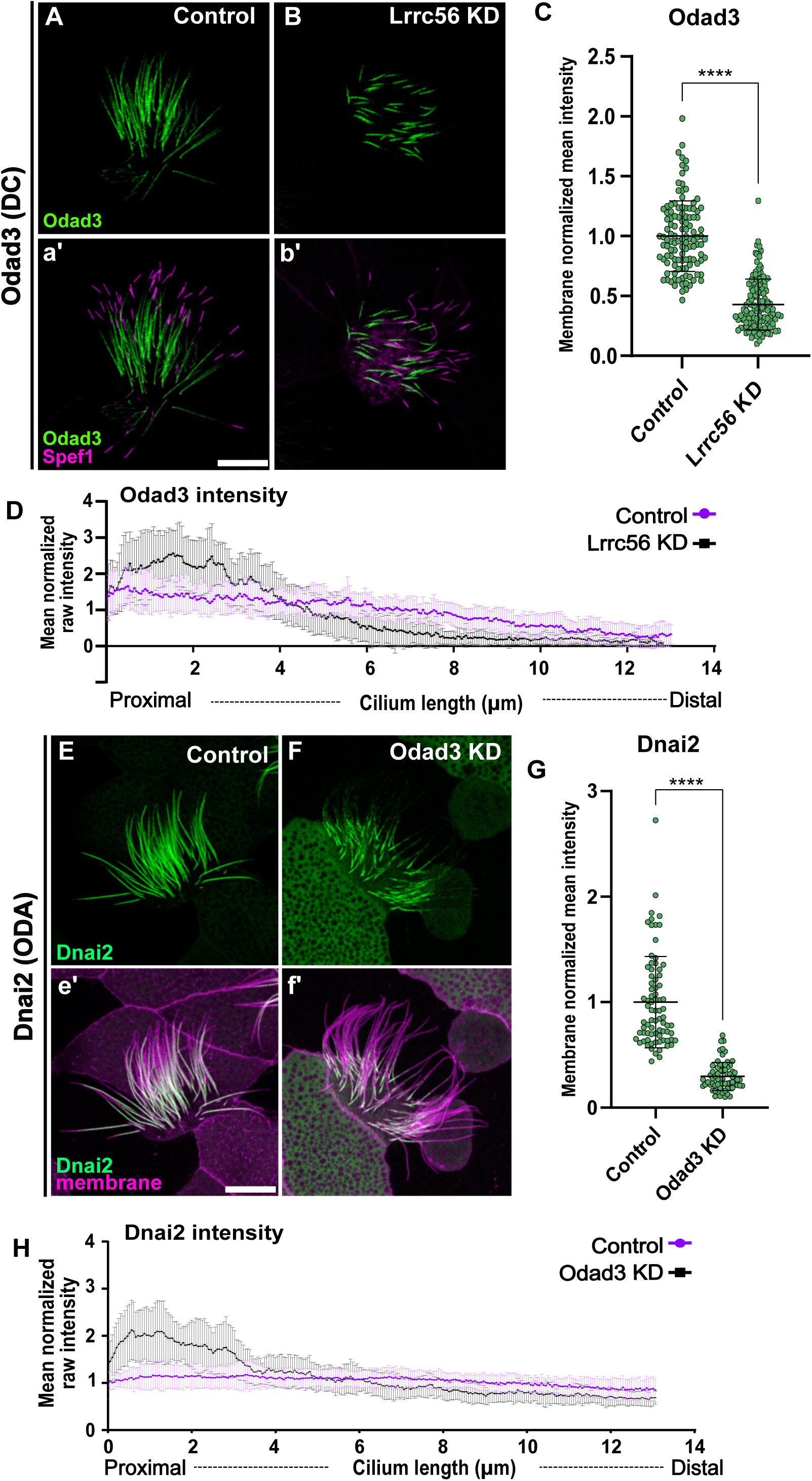
Lrrc56 knockdown alters Odad3 localization at distal end of the motile axoneme. (A) In control MCCs, the docking complex subunit Odad3 (green) localizes along the length of the motile cilium. (a′) Merged image showing Odad3 (green) and the distal tip marker Spef1 (magenta). Odad3 is enriched throughout the axoneme, excluding the Spef1-positive distal domain. (B) In Lrrc56 KD MCCs, Odad3 localization to axonemes is decreased. (b′) Merged image of Odad3 (green) and Spef1 (magenta) in Lrrc56 KD cilia shows reduced Odad3 intensity along the axoneme. (C) Quantification of normalized Odad3 mean intensity in control and Lrrc56 KD MCCs. Mean intensity was measured from the base of the cilium to the beginning of Spef1 domain in individual cilia, mean intensity was normalized to membrane marker, each dot represents a motile cilium. (D) Quantification of raw mean normalized Odad3 intensity along the axoneme in control and Lrrc56 KD MCCs. In Lrrc56 KD MCCs, Odad3 signal is enriched at the proximal end of the axoneme compared to controls. (Control: N = 33; Odad3 KD: N = 33). (E) The ODA subunit Dnai2 (green) localizes along the axoneme in control MCCs. (e′) Merged image of Dnai2 (green) and a membrane marker (magenta). (F) In Odad3 KD MCCs, Dnai2 intensity is reduced along the axoneme, particularly at the distal end. Quantification of Dnai2 mean intensity in control and Lrrc56 KD MCCs, normalized to a membrane marker. (H) Quantification of Dnai2 intensity along the cilium in control and *Odad3* KD MCCs. Plot shows representative axonemes (Control: N = 33; Odad3 KD: N = 21). **** P-value <0.0001. Scale bars = 10 µm.

Although the overall Odad3 signal was diminished, residual signal remained at the proximal region of the axoneme (Fig. 5B, b’). To further assess the distribution pattern, we quantified the raw Odad3-GFP intensity along the axoneme—excluding the Spef1-enriched distal domain—and normalized it to the mean Odad3-GFP intensity per cilium. In control cells, normalized Odad3-GFP intensity remained relatively constant along the first 8 µm of the axoneme (Fig. 5D). In contrast, in *lrrc56* morphants, Odad3 was strongly accumulated to above normal levels within the proximal 4 µm (Fig. 5D), suggesting that impaired distal deployment result to ectopic proximal Odad3.

The pentameric ODA-DC is essential for the proper assembly of ODAs along the microtubules (Owa et al., 2014; Takada et al., 2002). Thus, we examined the impact of Odad3 knockdown on the distribution of the ODA subunit Dnai2, which is typically distributed along the entire motile axoneme in control MCCs (Fig. 5D, d’). As expected, Odad3 depletion resulted in a substantial loss of ODA components in morphant MCCs (Fig. 5E, e’). Quantification of GFP-Dnai2 fluorescence mean intensity along the axoneme, normalized to a membrane marker, revealed a significant decrease in Dnai2 signal in *Odad3* knockdown cells compared to controls (Fig. 5F). Notably, despite overall loss, Dnai2 accumulated at the proximal axoneme, reminiscent of Odad3 behavior in Lrrc56-deficient MCCs (Fig. 5B, b’). To capture this shift, we measured raw GFP-Dnai2 intensity along the axoneme and normalized it to the mean intensity in each condition. As shown in Figure 5G, Dnai2 exhibited a proximal enrichment in Odad3-deficient cells.

### Odad3 variants of unknown function disrupt interaction with Lrrc56

A small number of validated PCD-causative variants in *CCDC151* (Odad3) have been reported in the literature (Alsaadi et al., 2014; Asseri et al., 2023; Hjeij et al., 2014). However, many ODAD3 variants of uncertain significance (VUS) are associated with ciliopathy in the human genetic database ClinVar. Notably, one such missense variant maps precisely to the Lrrc56-Odad3 interface (R207W) has been reported as a VUS in ClinVar.

This residue is well conserved in *Xenopus* (R171), so we examined its localization in MCCs. Compared to controls, the R171W variant of Odad3 displayed significantly reduced localization to the axoneme (Fig. 6 C, c’, D). Since this residue is predicted to be critical for interaction with Lrrc56, we assessed whether the R171W variant disrupts this interaction using co-immunoprecipitation (co-IP) assays. We co-immunoprecipitated Lrrc56 and probed for wild-type or R171W Odad3. As expected, wild-type Odad3 co-immunoprecipitated with LRRC56. However, the R171W variant showed reduced interaction with Lrrc56 compared to the wild-type protein (Fig. 6E). These findings demonstrate that a disease associated variant of Odad3 disrupts its interaction with Lrrc56, potentially leading to defects in ciliary structure or function.

**Figure 6.**
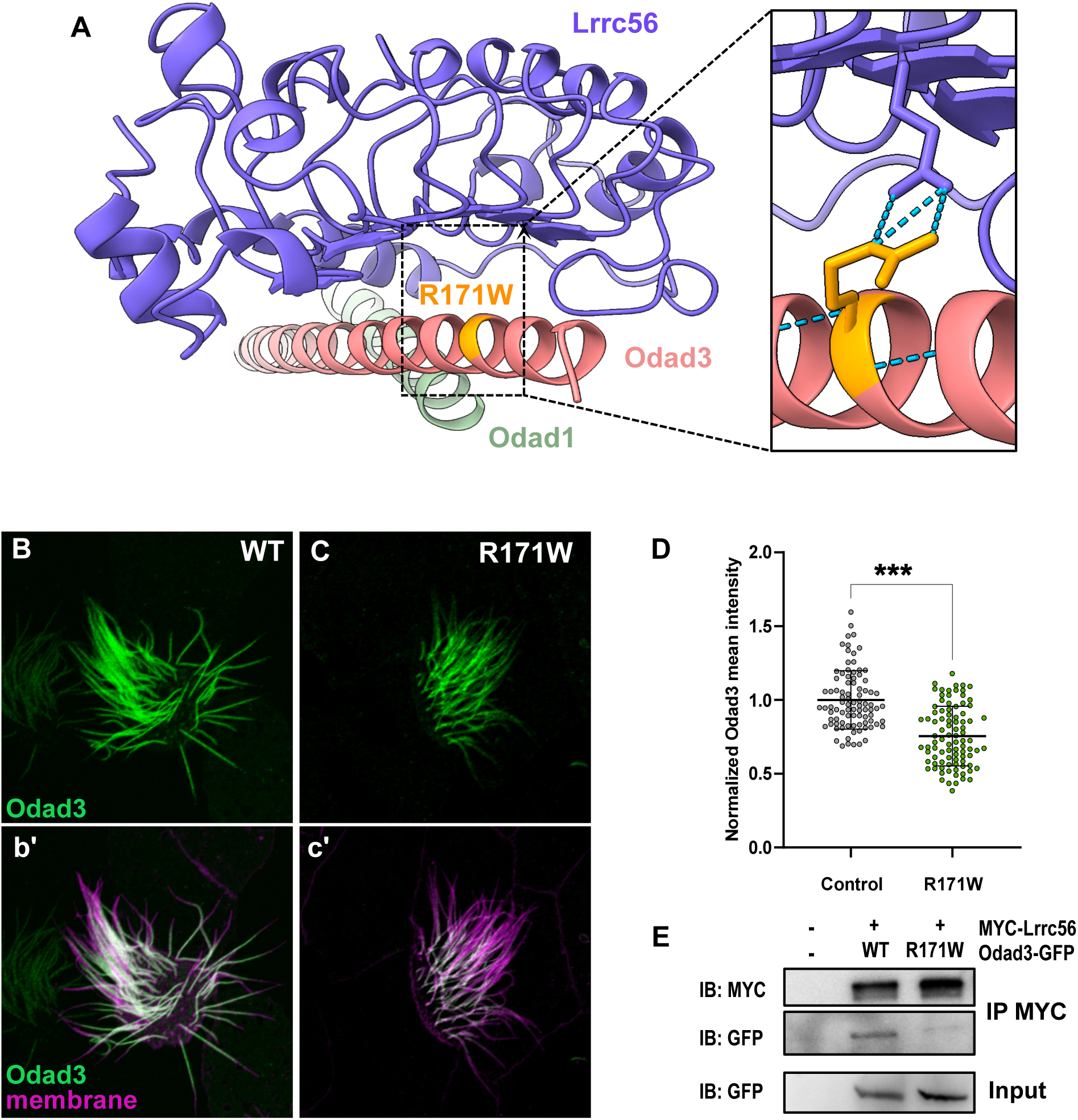
Odad3 variants of unknown function disrupt interaction with Lrrc56. (A) Alpha fold 3 model of *Xenopus* Lrrc56, Odad3, and Odad1 highlighting location of R171W (human E207W) variant of unknown function in Odad3. Close-up of the Lrrc56-Odad3 interface, showing predicted interaction of R171 residue and Lrrc56 E96 residue. (B) Wildtype localization of GFP-Odad3 in MCCs. Odad3 (green) localizes along the axoneme except for the most distal end of the cilium, motile cilia are labeled with membrane marker (magenta) (b’). (C) Expression of the R171W allele in MCCs disrupted its localization to axonemes compared to controls. (D) Quantification of normalized mean Odad3 intensity along the motile cilium showing decreased GFP signal R171W compared to wildtype control (WT). (E) Co-Immunoprecipitation (coIP) of Lrrc56 and Odad3. Representative Western blot of IP of the MYC-Lrrc56 and Odad3-GFP and of input GFP, between wildtype (WT) and R171W allele. *** P-value <0.0002. Scale bars = 10 µm.

## Discussion

*LRRC56* has recently emerged as a novel candidate gene implicated in PCD (Bonnefoy et al., 2018); but its functions have been explored largely in unicellular organisms. Here, we assessed the function of Lrrc56 specifically in vertebrates by leveraging the MCCs of *Xenopus* embryos and combining knockdown approaches with expression of human disease-associated variants. Our results reveal that Lrrc56 is required for the proper deployment of ODAs along MCC axonemes (Fig. 3B, F). Furthermore, we show that PCD-associated Lrrc56 variants impair the protein’s localization in a domain-specific manner (Fig. 2B–M), suggesting distinct structural contributions to Lrrc56 function. Together, our study provides new mechanistic insight into *LRRC56*-associated motile ciliopathies and highlights *Xenopus* as a powerful platform to investigate the function of poorly characterized ciliary genes *in vivo*.

Using affinity purification mass spectrometry, we identified Odad1 and Odad3—core components of the ODA docking complex—as candidate binding partners of Lrrc56. This interaction suggests that Lrrc56 may coordinate the transport or docking of ODAs in conjunction with ODA-DC components. Similar to observations in *Trypanosoma* (Bonnefoy et al., 2024), where *Lrrc56* mutants exhibit loss of distal docking complexes during flagellar elongation, we observed that loss of Lrrc56 in *Xenopus* MCCs causes distal depletion of ODAs and ODA-DC components, despite their accumulation at the proximal axoneme. This pattern was also observed in *Odad3* knockdown. Together, these data support a model in which Lrrc56 facilitates the distal trafficking or incorporation of ODA docking complex proteins, likely acting downstream of their initial assembly.

How pre-assembled ODA-DC are trafficked from their synthesis site to the cilium remains elusive. Recent work in *Tetrahymena* has shown that the conserved protein Shulin binds cytoplasmic ODAs, stabilizing them in a compact, transport-ready conformation (Mali et al., 2021). By analogy, Lrrc56 may similarly function in the cytosol to stabilize or “package” ODA docking complexes for delivery to the ciliary base. Within the axoneme, Lrrc56 could then act to promote region-specific trafficking or docking—consistent with its localization to DynAPs, basal bodies, and axonemes. The differential accumulation of ODAs and ODA docking complex components upon Lrrc56 loss suggests that vertebrate MCCs may possess mechanisms for spatial regulation of docking complex activity along the proximodistal axis.

Indeed, studies in *Chlamydomonas* and *Trypanosoma* have uncovered proximodistal patterning of ODA-DC components, with distinct proximal and distal docking modules that contribute to ciliary beat regulation (Dutcher, 2019). While mammalian ODA docking complex proteins CCDC114 and CCDC151 are reported to distribute repeatedly along the axoneme (Gui et al., 2021), our findings raise the possibility that subtle proximodistal asymmetries exist but remain unresolved in vertebrate systems. The compartmentalized localization of Lrrc56 and its role in coordinating distal ODA deployment may reflect such underlying specialization.

In summary, our work defines a conserved, vertebrate-specific role for Lrrc56 in ODA deployment and identifies key protein interactions disrupted in disease-linked alleles. By bridging insights from model organisms and human genetics, this study expands our understanding of motile ciliopathies and provides a framework for dissecting complex ciliary assembly pathways *in vivo*.

## Materials and Methods

### *Xenopus* embryo manipulations

All *Xenopus* experiments were conducted in accordance with the animal protocol AUP-2024-00130 and the animal ethics guidelines of the University of Texas at Austin. Female adult *Xenopus laevis* were induced to ovulate by injection of hCG (human chorionic gonadotropin). *In vitro* fertilization was carried out by homogenizing a small fraction of a testis in 1/3x Marc’s Modified Ringer’s (MMR). Embryos were dejellied in 1/3x MMR with 3%(w/v) cysteine at (pH7.8), microinjected with mRNA or morpholinos (MOs) in 2% ficoll (w/v) in 1/3x MMR. Injected embryos were washed with 1X MMR after at least 30 minutes and were incubated in 1/3x MMR until the appropriate stages.

### Plasmid, mRNA, and MO microinjections

*Xenopus* gene sequences were obtained from Xenbase (http://www.xenbase.org). Open reading frames (ORF) of genes were amplified from the *Xenopus laevis* cDNA library by polymerase chain reaction (PCR) and then inserted into a pCS10R vector containing a fluorescent tag. In addition to the vectors described previously (Huizar et al, 2018; Lee et al, 2020), the following constructs were cloned into pCS10R vector: GFP-Lrrc56, Flag-Lrrc56, MYC-Lrrc56, GFP-Odad1, Odad3-GFP. To generate a *Xenopus* allele corresponding to the human patient allele, mutagenesis was performed on the GFP-lrrc56 and GFP-odad3 plasmids, using Q5 Site-Directed Mutagenesis Kit (NEB, Cat #E0554S). All constructs were linearized and the capped mRNAs were synthesized using mMESSAGE mMACHINE SP6 transcription kit (ThermoFisher Scientific, Cat #AM1340). Each mRNA was injected into two ventral blastomeres at the four-cell stage (∼30-80 pg per blastomere). For APMS, GFP-lrrc56 and GFP-empty plasmids (35 pg per blastomere) were injected into both ventral and dorsal blastomeres at the four-cell stage. Morpholinos were designed to target 1^st^ or 3^rd^ exon-intron splicing junction of the alloallele of *lrrc56, odad3,* and *odad1* in the allotetraploid genome of *Xenopus laevis.* The morpholino sequences and the working concentrations include:

Lrrc56 MO: 5’ACTGAGTCTTAATGAAATCTTACCA 3’, 10-15 ng per injection

Odad3 MO: 5’CCTTTAATCAACTGACTTACCCAGG 3’, 10 ng per injection

### Live imaging and image analysis

Live imaging of multi-ciliated cells was performed in *Xenopus* embryos expressing tagged proteins at stages 24–26. Whole embryos were mounted between coverslips and immersed in 1/3X MMR prior to imaging. Imaging was conducted using a Zeiss LSM700 laser scanning confocal microscope equipped with a Plan-Apochromat 63×/1.4 NA oil immersion objective (Zeiss) or a Nikon Eclipse Ti confocal microscope with a 63×/1.4 NA oil immersion objective. Quantitative image analysis was performed using Fiji. Graphs were generated, and statistical analyses, including error bars representing mean ± SD and P values, were conducted using Prism 10 software. Statistical significance was determined using a non-parametric Mann-Whitney U test for two-group comparisons and a one-way ANOVA for comparisons involving more than two groups.

### Immunoprecipitation of *Xenopus* animal caps for mass-spectrometry

To identify Lrrc56 interactors, circular plasmids of GFP only, or GFP-lrrc56 driven by MCC-specific α-tubulin promoter were injected into 4-blastomeres of 4 cell stage *Xenopus* embryos. Approximately 550 animal caps per sample were isolated at stage eight using forceps and were cultured in 1X Steinberg’s solution (0.58 mM NaCl, 0.64 mM KCl, 0.33 mM Ca(NO2)2, 0.8 mM MgSO4, 5 mM Tris, 50 µg/ml gentamicin, pH 7.4–7.6) until sibling embryos reached stage 24. The cultured explants were collected, and immunoprecipitation (IP) was performed using GFP-Trap Agarose Kit (ChromoTek, cat# gtak-20). Immunoprecipitated proteins were eluted in 2X SDS-PAGE sample buffer.

### Affinity purification-mass spectrometry from JH 2025

Immunoprecipitated proteins in SDS-PAGE sample buffer were heated 5 min at 95°C before loading onto a 7.5% acrylamide mini-Protean TGX gel (BioRad). After 7 min of electrophoresis at 100 V the gel was stained with Imperial Protein stain (Thermo) according to manufacturer’s instructions. The protein band was excised, diced to 1 mm cubes and processed for in-gel trypsin digestion as in Goodman et al., 2018.

Digested peptides were desalted with 6µg-capacity ZipTips (Thermo Scientific), dried, and resuspended in 20µl of 5% acetonitrile, 0.1% acetic acid for mass-spectrometry. Peptides were separated using reverse phase chromatography on a Dionex Ultimate 3000 RSLCnano UHPLC system (Thermo Scientific) with a C18trap to EASY-Spray PepMap RSLC C18 column (Thermo Scientific, ES902) configuration eluted with a 3% to 45% gradient over 60 min. Spectra were collected on a Thermo Orbitrap Fusion Lumos Tribrid mass spectrometer using a data-dependent top speed HCD acquisition method with full precursor ion scans (MS1) collected at 120,000 m/z resolution. Monoisotopic precursor selection and charge-state screening were enabled using Advanced Peak Determination (APD), with ions of charge + two selected for high energy-induced dissociation (HCD) with stepped collision energy of 30% +/- 3% Dynamic exclusion was active for ions selected once with an exclusion period of 20 s. All MS2 scans were centroid and collected in rapid mode. Raw MS/MS spectra were processed using Proteome Discoverer (v2.5) and the Percolator node to assign unique peptide spectral matches (PSMs) and protein assignments (FDR .01) to a *X. laevis* proteome derived from the 2023 UniProt *Xenopus laevis* reference proteome of 35,860 protein sequences with homeologs and highly related entries collapsed into EggNOG vertebrate-level orthology groups (Huerta-Cepas et al., 2016). This database and the mass spectrometry data are available on MassIVE dataset MSV000098174 (https://massive.ucsd.edu/ProteoSAFe/static/massive.jsp).

In order to identify proteins significantly associated with each bait, we used the degust statistical framework (https://degust.erc.monash.edu/) to calculate both a log2 fold-change and an FDR for each protein enrichment based on the observed PSMs in the bait versus control pulldown. Settings used were “RUV (edgeR-quasi-likelihood), Normalization TMM, and Flavour RUVr” and at least 2 counts in at least 2 samples.

### RT-PCR

To confirm the efficacy of Lrrc56 and Odad3 morpholinos, morpholinos were injected into all cells at 4-cell stage of embryo. Total RNA was isolated with Trizol reagent (Invitrogen cat # 15596026) at stage 26, and cDNA was synthesized with M-MLV reverse transcription kit (Invitrogen cat# 28025013). Lrrc56, Odad3 and Odc1 were amplified by Taq-polymerase (Invitrogen cat# 10342020) with the following primers: Lrrc56.L 55F 5’ gatttgggttggcaaggatta 3’, Lrrc56.L 508R 5’ ctccagattgtttccctcaaga 3’, Odad3.L MO 63F 5’ gcacaagaaactccagctcc 3’, and Odad3.L MO 367R 5’ tcttcttctctgcctggtgg 3’.

### Immunoblotting

Embryos were lysed in Lysis buffer (ChromoTek, cat# gtak-20) supplemented with protease inhibitors. The lysates were centrifuged to remove cell debris, and the supernatants were subjected to SDS-PAGE followed by immunoblotting using standard protocols. The antibodies used were as follows: anti-GFP antibody (Santa Cruz, Cat# sc-9996), HRP-conjugated goat anti mouse IgG (H+L) secondary antibody (ThermoFisher Scientific, Cat #31430), beta actin monoclonal antibody (Proteintech, Cat #6009-1), anti-Myc antibody (Abcam, Cat #9E10).

## Acknowledgements

We thank Xenbase and the European Xenopus Resource Centre (EXRC) for providing *Xenopus* resources, and Tynan Gardner for generating the initial AlphaFold models.

## Competing interests

Authors declare no competing of financial interests.

## Funding

JBW is supported by the NHLBI (R01HL117164); NGR-N was supported by a Provost’s Early Career Fellowship from the University of Texas at Austin. JH was supported by the Korea Health Technology R&D Project through Korea Health Industry Development Institute (KHIDI), funded by the Ministry of Health & Welfare, Republic of Korea (grant number: HI19C1095). EMM was supported by the NIGMS (R35GM122480), Army Research Office (W911NF-12-1-0390), and the Welch Foundation (F-1515).

## Author contributions statement

Conceptualization: NGRN., CJL., JBW; Data curation: NGRN., CJL., JBW; Formal analysis: NGRN., CJL., OP.; Funding acquisition: JBW, EMM; Investigation: NGRN., CJL., JH., OP. Methodology: NGRN., CJL., JBW; Project administration: NGRN., JBW., Resources: JBW., EMM., Supervision: NGRN., JBW, EMM; Writing – original draft: NGRN., JBW.; Writing – review & editing: NGRN., JBW.

## Data and resource availability

The database and the mass spectrometry data here reported are available as MassIVE dataset MSV000098174 (https://massive.ucsd.edu/ProteoSAFe/static/massive.jsp).

## Supplementary Table Legend

**Supplementary Table 1. Table showing orthogroups and proteins with PSMs identified by APMS with Lrrc56.**

---

**Supplemental Figure 1.**
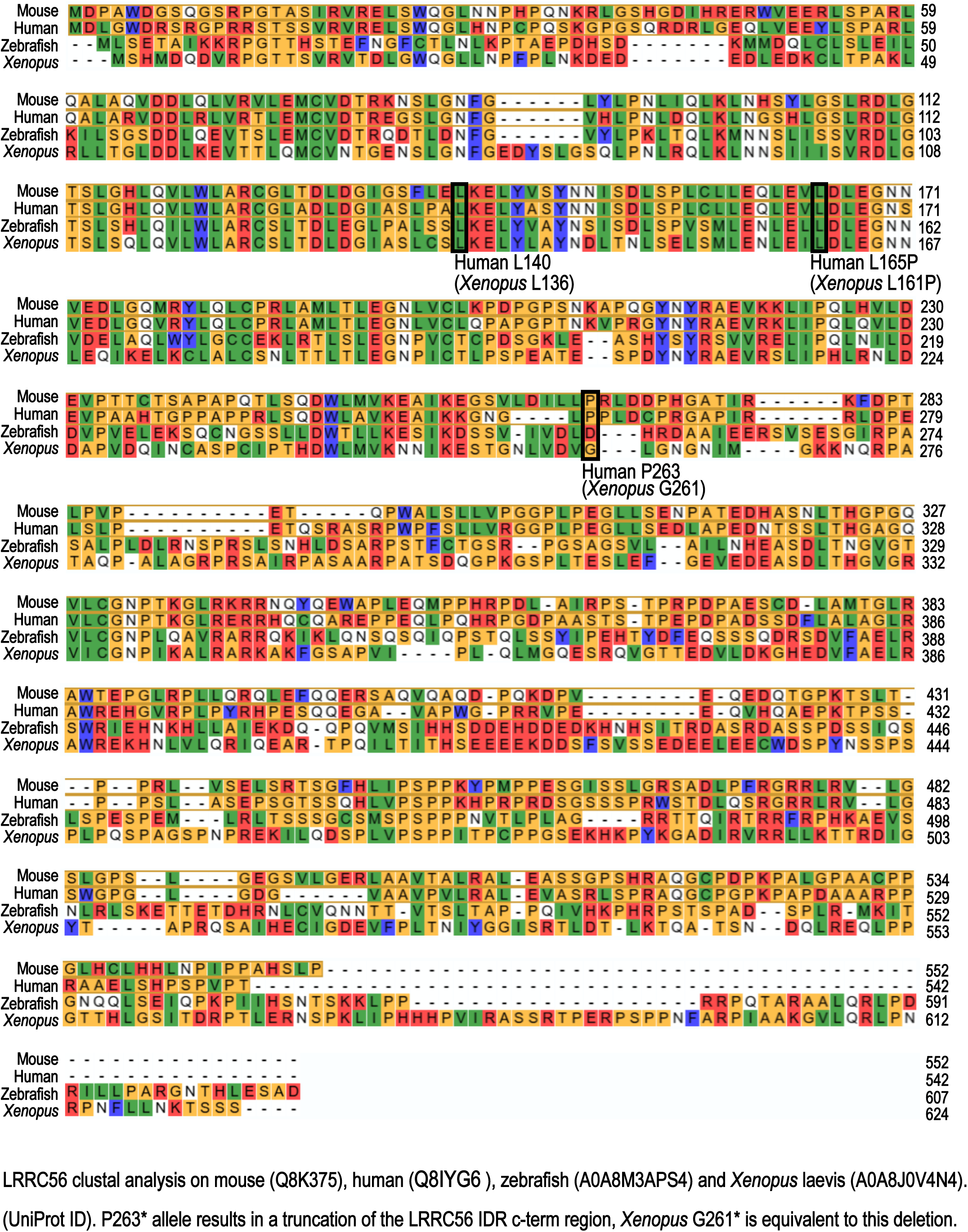
Alignment of vertebrate Lrrc56 proteins. Multiple sequence alignment showing conserved Lrrc56 residues across mouse, human, zebrafish, and *Xenopus.* Rectangles indicate conserved ciliopathy loci characterized here.

**Supplemental Figure 2.**
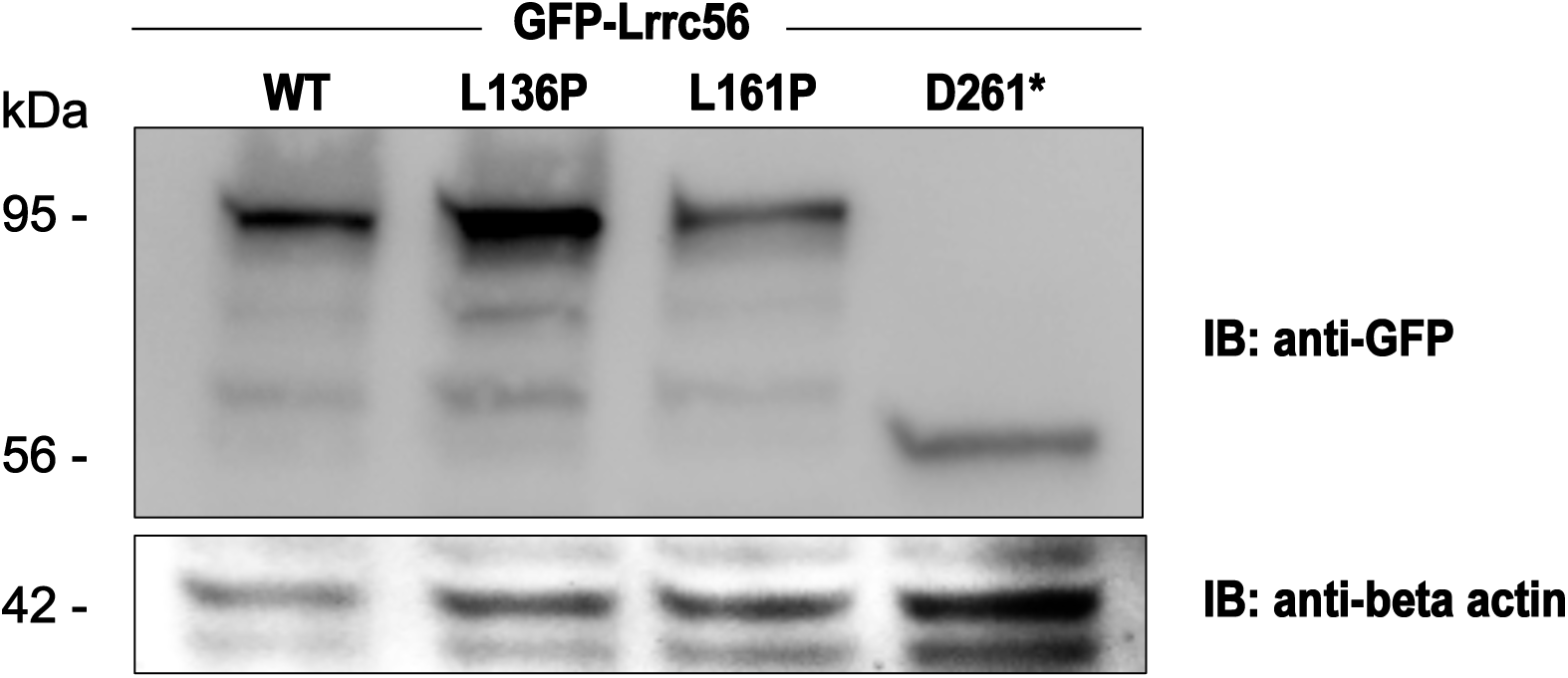
Lrrc56 ciliopathy variants protein abundance. Western blot showing protein levels for indicated disease alleles when expressed in *Xenopus* embryos. *Xenopus* variants (human): L136P (L140P), L161P (L165P), G261* (P263*). Western blot of total protein from N=20 embryos NF 25, injected with 80pg of GFP-Lrrc56 WT, L136P, L161P and 160pg of D261*. Anti GFP 1:200 and Anti B-actin housekeeping control (1:10,000).

**Supplemental Figure 3.**
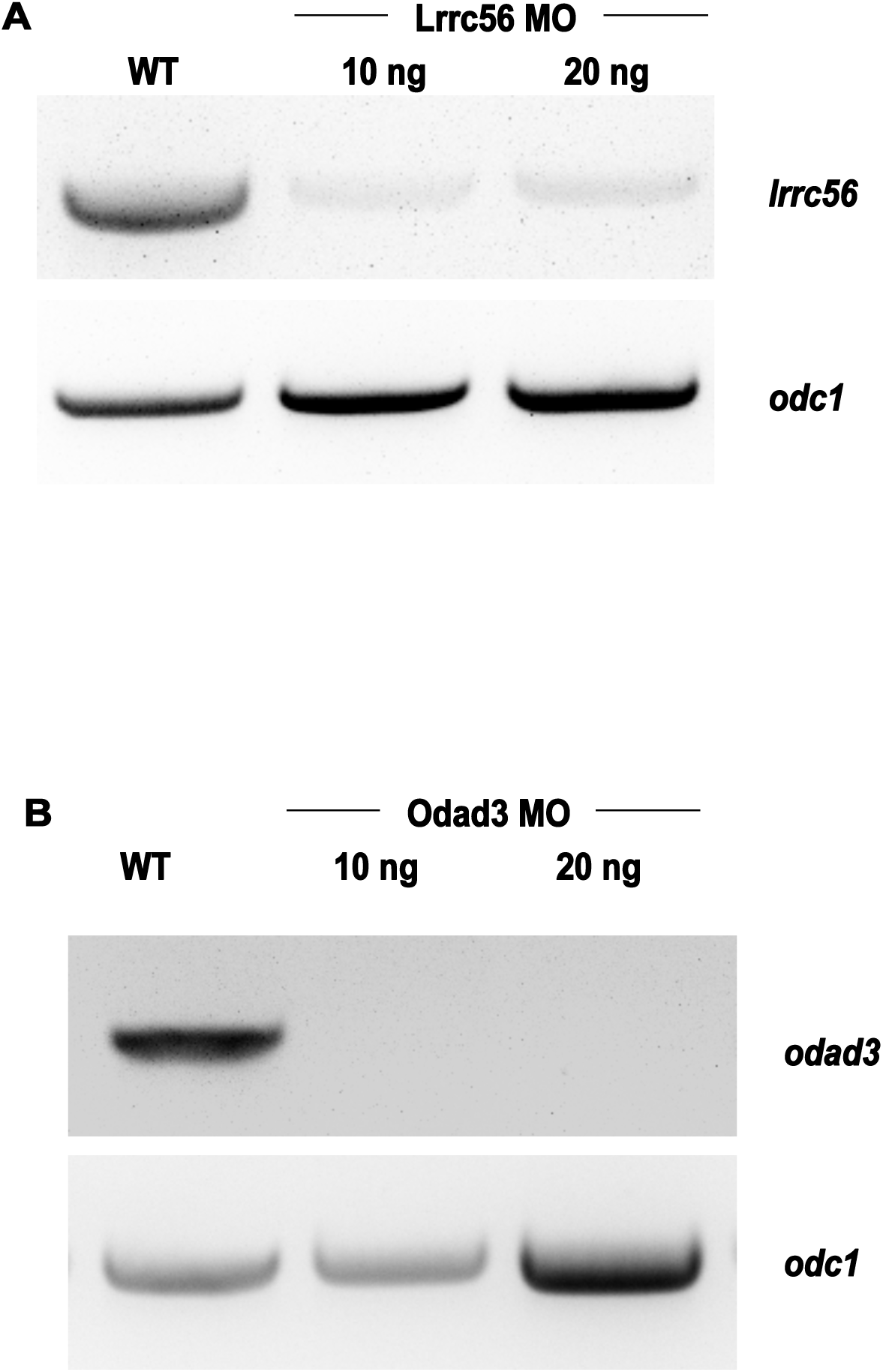
Validation of Lrrc56 and Odad3 splice-blocking morpholino efficiency by RT-PCR. (A) Gel image of RT-PCR of *lrrc56* and *odc1* mRNA levels in wildtype control (WT), Lrrc56 MO 10ng and 20ng injected embryos. (B) Gel image of RT-PCR of *odad3* and *odc1* mRNA levels in wildtype control (WT), Lrrc56 MO 10ng and 20ng injected embryos.

**Supplemental Figure 4.**
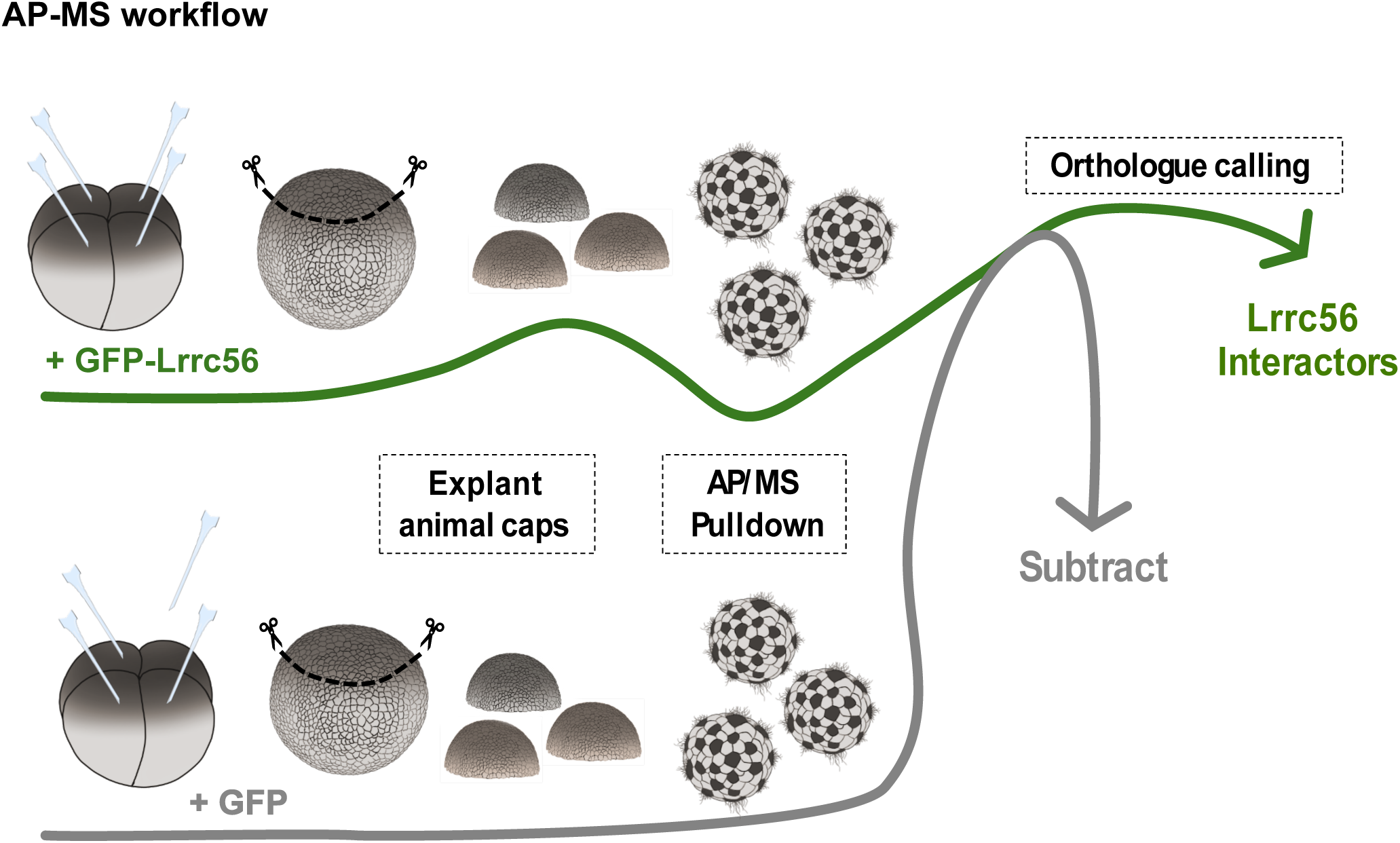
Schematic of AP-MS workflow for identification of *in vivo* Lrrc56 interactors. A plasmid encoding GFP-tagged Lrrc56 under the control of the MCC-specific α-tubulin promoter was injected into Xenopus embryos at the 2–4 cell stage (stage 3). Animal cap explants were dissected at stage 8 and cultured until the early stage of ciliogenesis (stage 23). Explants were then harvested and subjected to GFP-based immunoprecipitation followed by affinity purification mass spectrometry (AP-MS). A parallel experiment using unfused GFP was performed to account for non-specific interactions, and these were subtracted from the experimental dataset to identify specific Lrrc56 interactors.

**Supplemental Figure 5.**
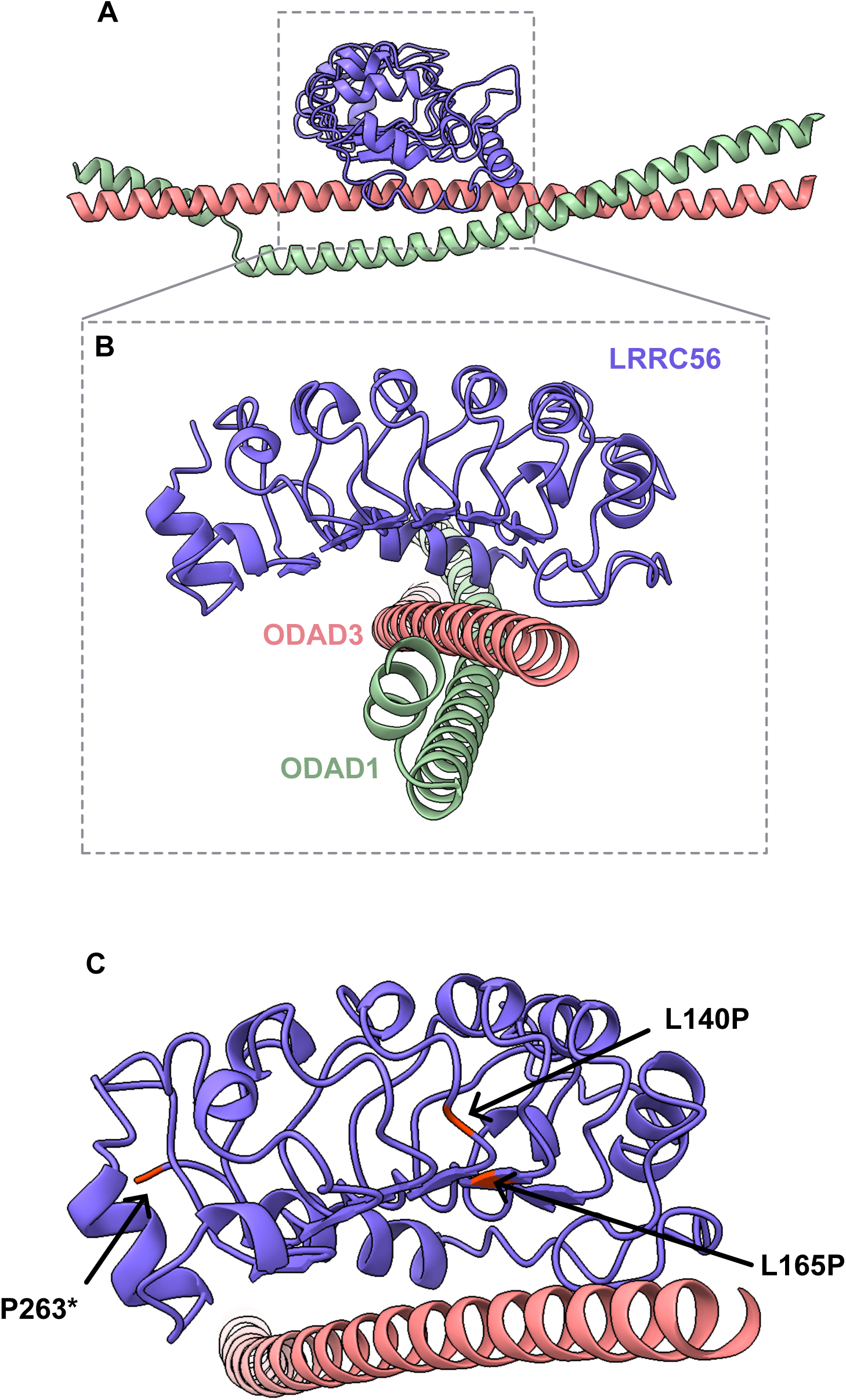
Human AF3 model of LRRC56, ODAD3 and ODAD1. (A) AlphaFold3-predicted structure of human LRRC56 and its interactors, ODAD3 and ODAD1. Each monomer is color-coded as indicated. The model includes residues 53-263 of LRRC56, 159-315 of ODAD3, and 80-224 of ODAD1. (B) Enlarged view of the LRRC56–ODAD3 interface (C) Close-up of the LRRC56–ODAD3 interface in the AlphaFold3 model, highlighting the positions of Lrrc56 ciliopathy-associated variants (L140P, L165P, and P263*).

## Notes

### Competing Interest Statement

The authors have declared no competing interest.

## References

1. A. Deblandre, G., Wettstein, D. A., Koyano-Nakagawa, N., & Kintner, C. (1999). A two-step mechanism generates the spacing pattern of the ciliated cells in the skin of Xenopus embryos. Development, 126(21), 4715–4728. 10.1242/dev.126.21.4715

2. Alasmari, B. G., Saeed, M., Alomari, M. A., Alsumaili, M., & Tahir, A. M. (2022). Primary Ciliary Dyskinesia: Phenotype Resulting From a Novel Variant of LRRC56 Gene. Cureus, 14(8), e28472. 10.7759/cureus.28472

3. Alsaadi, M. M., Erzurumluoglu, A. M., Rodriguez, S., Guthrie, P. A. I., Gaunt, T. R., Omar, H. Z., Mubarak, M., Alharbi, K. K., Al-Rikabi, A. C., & Day, I. N. M. (2014). Nonsense Mutation in Coiled-Coil Domain Containing 151 Gene (CCDC151) Causes Primary Ciliary Dyskinesia. Human Mutation, 35(12), 1446–1448. 10.1002/humu.22698

4. Asseri, A. A., Shati, A. A., Asiri, I. A., Aldosari, R. H., Al-Amri, H. A., Alshahrani, M., Al-Asmari, B. G., & Alalkami, H. (2023). Clinical and Genetic Characterization of Patients with Primary Ciliary Dyskinesia in Southwest Saudi Arabia: A Cross Sectional Study. Children, 10(10), 1684. 10.3390/children10101684

5. Bonnefoy, S., Alves, A. A., Bertiaux, E., & Bastin, P. (2024). LRRC56 is an IFT cargo required for assembly of the distal dynein docking complex in Trypanosoma brucei. Molecular Biology of the Cell, 35(8), ar106. 10.1091/mbc.E23-11-0425

6. Bonnefoy, S., Watson, C. M., Kernohan, K. D., Lemos, M., Hutchinson, S., Poulter, J. A., Crinnion, L. A., Berry, I., Simmonds, J., Vasudevan, P., O’Callaghan, C., Hirst, R. A., Rutman, A., Huang, L., Hartley, T., Grynspan, D., Moya, E., Li, C., Carr, I. M., … Sheridan, E. G. (2018). Biallelic Mutations in LRRC56, Encoding a Protein Associated with Intraflagellar Transport, Cause Mucociliary Clearance and Laterality Defects. American Journal of Human Genetics, 103(5), 727–739. 10.1016/j.ajhg.2018.10.003

7. Desai, P. B., Freshour, J. R., & Mitchell, D. R. (2015). Chlamydomonas Axonemal Dynein Assembly Locus ODA8 Encodes a Conserved Flagellar Protein Needed for Cytoplasmic Maturation of Outer Dynein Arm Complexes. *Cytoskeleton (Hoboken*, N.j.), 72(1), 16–28. 10.1002/cm.21206

8. Drysdale, T. A., & Elinson, R. P. (1992). Cell Migration and Induction in the Development of the Surface Ectodermal Pattern of the Xenopus laevis Tadpole. *Development*, Growth & Differentiation, 34(1), 51–59. 10.1111/j.1440-169X.1992.00051.x

9. Fowkes, M. E., & Mitchell, D. R. (1998). The Role of Preassembled Cytoplasmic Complexes in Assembly of Flagellar Dynein Subunits. Molecular Biology of the Cell, 9(9), 2337–2347. 10.1091/mbc.9.9.2337

10. Goodman, J. K., Zampronio, C. G., Jones, A. M. E., & Hernandez-Fernaud, J. R. (2018). Updates of the In-Gel Digestion Method for Protein Analysis by Mass Spectrometry. Proteomics, 18(23), 1800236. 10.1002/pmic.201800236

11. Gui, M., Farley, H., Anujan, P., Anderson, J. R., Maxwell, D. W., Whitchurch, J. B., Botsch, J. J., Qiu, T., Meleppattu, S., Singh, S. K., Zhang, Q., Thompson, J., Lucas, J. S., Bingle, C. D., Norris, D. P., Roy, S., & Brown, A. (2021). *De novo* identification of mammalian ciliary motility proteins using cryo-EM. Cell, 184(23), 5791–5806.e19. 10.1016/j.cell.2021.10.007

12. Hibbard, J. V. K., Vázquez, N., & Wallingford, J. B. (2022). Cilia proteins getting to work – how do they commute from the cytoplasm to the base of cilia? Journal of Cell Science, 135(17), jcs259444. 10.1242/jcs.259444

13. Hjeij, R., Onoufriadis, A., Watson, C. M., Slagle, C. E., Klena, N. T., Dougherty, G. W., Kurkowiak, M., Loges, N. T., Diggle, C. P., Morante, N. F. C., Gabriel, G. C., Lemke, K. L., Li, Y., Pennekamp, P., Menchen, T., Konert, F., Marthin, J. K., Mans, D. A., Letteboer, S. J. F., … Mitchison, H. M. (2014). *CCDC151* Mutations Cause Primary Ciliary Dyskinesia by Disruption of the Outer Dynein Arm Docking Complex Formation. The American Journal of Human Genetics, 95(3), 257–274. 10.1016/j.ajhg.2014.08.005

14. Horani, A., & Ferkol, T. W. (2021). Understanding primary ciliary dyskinesia and other ciliopathies. The Journal of Pediatrics, 230, 15–22.e1. 10.1016/j.jpeds.2020.11.040

15. Huizar, R. L., Lee, C., Boulgakov, A. A., Horani, A., Tu, F., Marcotte, E. M., Brody, S. L., & Wallingford, J. B. (2018). A liquid-like organelle at the root of motile ciliopathy. eLife, 7, e38497. 10.7554/eLife.38497

16. Kamiya, R. (1988). Mutations at Twelve Independent Loci Result in Absence of Outer Dynein Arms in Chflamydomonasreinhardtii. The Journal of Cell Biology, 107.

17. Knowles, M. R., Leigh, M. W., Ostrowski, L. E., Huang, L., Carson, J. L., Hazucha, M. J., Yin, W., Berg, J. S., Davis, S. D., Dell, S. D., Ferkol, T. W., Rosenfeld, M., Sagel, S. D., Milla, C. E., Olivier, K. N., Turner, E. H., Lewis, A. P., Bamshad, M. J., Nickerson, D. A., … Zariwala, M. A. (2013). Exome Sequencing Identifies Mutations in *CCDC114* as a Cause of Primary Ciliary Dyskinesia. The American Journal of Human Genetics, 92(1), 99–106. 10.1016/j.ajhg.2012.11.003

18. Lechtreck, K. (2022). Cargo adapters expand the transport range of intraflagellar transport. Journal of Cell Science, 135(24), jcs260408. 10.1242/jcs.260408

19. Lee, C., Cox, R. M., Papoulas, O., Horani, A., Drew, K., Devitt, C. C., Brody, S. L., Marcotte, E. M., & Wallingford, J. B. (2020). Functional partitioning of a liquid-like organelle during assembly of axonemal dyneins. eLife, 9, e58662. 10.7554/eLife.58662

20. Leslie, J. S., Hjeij, R., Vivante, A., Bearce, E. A., Dyer, L., Wang, J., Rawlins, L., Kennedy, J., Ubeyratna, N., Fasham, J., Irons, Z. H., Craig, S. B., Koenig, J., George, S., Pode-Shakked, B., Bolkier, Y., Barel, O., Mane, S., Frederiksen, K. K., … Baple, E. L. (2022). Biallelic DAW1 variants cause a motile ciliopathy characterized by laterality defects and subtle ciliary beating abnormalities. Genetics in Medicine : Official Journal of the American College of Medical Genetics, 24(11), 2249–2261. 10.1016/j.gim.2022.07.019

21. Li, P., He, Y., Cai, G., Xiao, F., Yang, J., Li, Q., & Chen, X. (2019). CCDC114 is mutated in patient with a complex phenotype combining primary ciliary dyskinesia, sensorineural deafness, and renal disease. Journal of Human Genetics, 64(1), 39–48. 10.1038/s10038-018-0514-z

22. Mali, G. R., Ali, F. A., Lau, C. K., Begum, F., Boulanger, J., Howe, J. D., Chen, Z. A., Rappsilber, J., Skehel, M., & Carter, A. P. (2021). Shulin packages axonemal outer dynein arms for ciliary targeting. *Science (New York*, N.Y.), 371(6532), 910–916. 10.1126/science.abe0526

23. Owa, M., Furuta, A., Usukura, J., Arisaka, F., King, S. M., Witman, G. B., Kamiya, R., & Wakabayashi, K. (2014). Cooperative binding of the outer arm-docking complex underlies the regular arrangement of outer arm dynein in the axoneme. Proceedings of the National Academy of Sciences of the United States of America, 111(26), 9461–9466. 10.1073/pnas.1403101111

24. Qiu, T., & Roy, S. (2022). Ciliary dynein arms: Cytoplasmic preassembly, intraflagellar transport, and axonemal docking. Journal of Cellular Physiology, 237(6), 2644–2653. 10.1002/jcp.30689

25. Raidt, J., Loges, N. T., Olbrich, H., Wallmeier, J., Pennekamp, P., & Omran, H. (2023). Primary ciliary dyskinesia. La Presse Médicale, 52(3), 104171. 10.1016/j.lpm.2023.104171

26. Takada, S., Wilkerson, C. G., Wakabayashi, K., Kamiya, R., & Witman, G. B. (2002). The Outer Dynein Arm-Docking Complex: Composition and Characterization of a Subunit (Oda1) Necessary for Outer Arm Assembly. Molecular Biology of the Cell, 13(3), 1015–1029. 10.1091/mbc.01-04-0201

27. Walentek, P., & Quigley, I. K. (2017). What we can learn from a tadpole about ciliopathies and airway diseases – Using systems biology in Xenopus to study cilia and mucociliary epithelia. Genesis (New York, N.Y. : 2000), 55(1–2), 10.1002/dvg.23001. https://doi.org/10.1002/dvg.23001

28. Wallmeier, J., Nielsen, K. G., Kuehni, C. E., Lucas, J. S., Leigh, M. W., Zariwala, M. A., & Omran, H. (2020). Motile ciliopathies. Nature Reviews Disease Primers, 6(1), Article 1. 10.1038/s41572-020-0209-6

29. Wu, R., Li, H., Wu, P., Yang, Q., Wan, X., & Wu, Y. (2025). LRRC56 deletion causes primary ciliary dyskinesia in mice characterized by dynein arms defects. Biology Open, 14(2), bio061846. 10.1242/bio.061846

30. Yamaguchi, H., Morikawa, M., & Kikkawa, M. (2023). Calaxin stabilizes the docking of outer arm dyneins onto ciliary doublet microtubule in vertebrates. eLife, 12, e84860. 10.7554/eLife.84860

